# The improved auxin signalling via *entire* mutation enhances aluminium tolerance in tomato

**DOI:** 10.1101/2025.08.29.673006

**Authors:** Regiane K. G. Silva, João Antonio Siqueira, Willian Batista-Silva, Marcelle Ferreira Silva, Thiago Wakin, Jonas Rafael Vargas, Gabriel Vilela, Robson Ribeiro, Domingos F.M. Neto, Aristea A. Azevedo, Cleberson Ribeiro, Alisdair R. Fernie, Adriano Nunes-Nesi, Wagner L. Araújo

## Abstract

Acidic soils limit food production in many developing countries by promoting the solubilization of aluminium (Al) cations. Consequently, roots absorb this metal from soil solution, arresting their growth while reducing water and nutrient uptake. To mitigate the impacts of Al, plants rewire their metabolism and growth, with diverse mechanisms in this process shaped by changes in auxin signalling. Here, we present a comprehensive study on the significance of auxin signalling for Al tolerance. We used tomato mutants with reduced (*diageotropica*, *dgt*) and increased (*entire*) auxin sensitivity to assess the regulation of growth and metabolism in plants coping with Al toxicity. Our results indicated a reduced Al tolerance in the *dgt* mutant, whereas *entire* was able to tolerate toxic levels of Al. This contrast can be explained by a differential accumulation of reactive oxygen species (ROS) in the root transition zone, where *dgt* exhibited more differentiated cells, making the pericycle evident. In contrast, *entire* showed only slight alterations in the transition zone, with root meristematic cells maintaining a reduced level of cell differentiation, which can be associated with sustained growth under toxic Al levels. These differences were followed by alterations in metabolites related to Al sensitivity in the roots of *dgt* plants, whereas the *entire* mutant exhibited only slight metabolic changes. Collectively, our results suggest that genetic modifications to regulate auxin perception have the potential to increase Al tolerance in crops.

## 1. INTRODUCTION

Throughout human history, the establishment of civilizations has primarily occurred in regions with fertile soils that enable food production. However, with the intense use of these soils and significant population growth, new civilizations have emerged in regions with low-fertility soils. This shift was driven, among other factors, by the ability to correct soil acidity using limestone. In acidic soils (pH ≤ 5.5) without limestone application, aluminium cations (mainly Al^3+^) are commonly solubilized (Kochian et al., 2004). Even after centuries, Al continues to limit soil fertility, forcing farmers to apply large amounts of chemical fertilizers, which poses a barrier to food production in developing countries. Plant roots are more affected by Al stress than shoots due to their direct contact with the metal, which inhibits water and nutrient uptake (Kochian et al., 2004; Kochian et al., 2015; Siqueira et al., 2022a, 2022b). Therefore, understanding the mechanisms that allow root growth under Al toxicity is fundamental to developing crops capable of maintaining productivity in soils with high acidity.

Despite growing knowledge of plant responses to Al toxicity, the most well-characterized mechanism promoting Al tolerance involves organic acids (OAs) (Nunes-Nesi et al., 2014; Kochian et al., 2015; Siqueira et al., 2022a). The exudation of OAs from root cells into the rhizosphere, where OAs bind to Al, neutralizes this metal. These molecules facilitate the sequestration of Al into vacuoles or can be exuded from root tip cells into the rhizosphere to prevent the absorption of Al ions (Delhaize et al., 2007; Kochian et al., 2015). OAs play critical roles in diverse pathways associated with plant growth and development. The physiological relevance of these metabolites varies across species and even among genotypes within the same species, where similar alterations in OA levels can result in varying degrees of Al tolerance (Zhang and Fernie, 2018; Pereira-Lima et al., 2023). It is essential to investigate how growth-promoting mechanisms are affected under Al stress conditions and whether inherent pathways interact with metabolic changes triggered by Al stress.

The initial damage occurs in the apoplast, where cell elongation is inhibited due to Al binding to the cell walls of the rhizodermis. This process prevents the cell wall remodelling necessary for cell elongation, a process mediated by phytohormones (Kopittke et al., 2015; Yang et al., 2017). Al is known to inhibit root growth by interacting with auxin signalling (Kollmeier et al., 2000; Kopittke, 2016; Zhang et al., 2018). Elevated auxin levels have been associated with Al sensitivity, which is correlated with the repression of the *Aluminium Sensitive 1* (*ALS1*) gene. This gene encodes a transporter responsible for reallocating Al into vacuoles, regulating Al detoxification in the symplast (Zhu et al., 2013). Consequently, exogenous application of auxin to the roots of maize (*Zea mays*) and alfalfa (*Medicago sativa*) has been shown to mitigate the impact of Al in inhibiting growth (Kollmeier et al., 2000; Wang et al., 2016a). Furthermore, auxin enhances Al-induced citrate exudation in soybean (*Glycine max*) roots by inducing the expression of *MATE* (*Multidrug and toxic compound extrusion*) genes (Wang et al., 2016b). Enhanced auxin signalling under Al toxicity appears to block auxin efflux into root tips, thereby improving Al tolerance (Zhang et al., 2018). Since differential auxin signalling can shape the growth and metabolism of OAs (Batista-Silva et al., 2019), it seems important to determine the extent to which auxin signalling shapes the anatomy and physiology of plants under Al stress.

Here, we present a comprehensive study on the significance of auxin signalling for Al tolerance. To this end, we used tomato (*Solanum lycopersicum*) cv. Micro-Tom mutants with reduced (*diageotropica*, *dgt*) and increased (*entire*) auxin sensitivity to assess traits relevant to Al stress responses. The mutant *dgt* is auxin-resistant whereas the levels of auxin are not altered by the mutation, whereas this mutation affects the transport of the hormone in roots (Balbi and Lomax, 2003; Mignolli et al., 2012; Ivanchenko et al., 2015). The *entire* mutant is hypersensitive to auxin exhibiting stronger responses to auxin application, wherein the maximum hypocotyl elongation of *entire* was reached with an auxin concentration 10-fold lower than wild-type (WT) plants (Wang et al., 2005). Consequently, these mutants encompass a valuable tool to understand the interactions between auxin sensitiveness and Al tolerance. Our results indicate that alterations in auxin signalling modulate crucial responses to Al stress, influencing not only morphophysiological traits but also shifts in the root elongation zone. We demonstrate that reduced auxin perception in *dgt* results in sensitivity to Al stress, whereas enhanced perception in *entire* promotes Al tolerance. These responses are predominantly mediated by changes in central metabolism, including metabolites such as sugars, proteins, and amino acids. Therefore, modulating auxin signalling pathways offers a promising strategy for enhancing crop performance under Al stress conditions.

## 2. MATERIAL AND METHODS

### 2.1 Plant material and growth phenotypes

Tomato (*Solanum lycopersicum* cultivar Micro-Tom) seeds of the wild type (WT) and lines with low auxin perception (*diageotropica, dgt*) and increased auxin signalling (*entire*, *AUX/IAA9*) in the Micro-Tom genetic background were obtained as previously described (Carvalho et al., 2011). The *dgt* mutation was obtained through EMS (Ethyl methanesulfonate) creating a stop codon at gene cyclophilin, whereas the EMS modified the gene *IAA9* generating a truncated polypeptide (Wang et al., 2005; Oh et al., 2006; Ivanchenko et al., 2015). The phenotypes of the *dgt* and *entire* plants in the Micro-Tom genetic background resemble those previously published for the same mutations in other tomato cultivars (Wang et al., 2005; Oh et al., 2006; Ivanchenko et al., 2015). The seeds were disinfected in a solution containing 0.5% (w/v) sodium hypochlorite for 5 minutes, followed by three washes with distilled water. They were then sown in “Germitest” paper rolls, which were moistened daily with distilled water. The seeds were kept in darkness for 8 days, and then exposed to light for 3 days before hydroponics experiments were initiated. The plants were cultivated in growth chambers under controlled conditions: a temperature of 25 °C, a 16 h light/8 h dark photoperiod, and light intensity of 330 µmol photons m^-2^ s^-1^.

Two hydroponic experiments with different exposure times to AlCl3 were implemented. In the first experiment, young plants with the first pair of true leaves (approximately 10 days after germination) were transferred to plastic pots containing a modified Hoagland and Arnon (1950) nutrient solution. This solution specifically contained 500 mmol L^−1^ Ca(NO3)2.4H2O, 500 mmol L^−1^ KNO3, 100 mmol L^−1^ MgSO4.7H2O, 50 mmol L^−1^ KH2PO4, 47.8 mmol L^−1^ H3BO3, 14.38 mmol L^−1^ MnCl2.4H2O, 1.36 mmol L^−1^ ZnSO4.7H2O, mmol L^−1^ CuSO4.5H2O, 0.12 mmol L^−1^ H2MoO4.H2O, and 420 µmmol L^−1^ of Fe-EDTA, adjusted to pH 4.5. The seedlings were maintained in an aerated plastic pot system with 4 seedlings per box. After 3 days of acclimation, the seedlings were transferred to a new nutrient solution with the same composition but subjected to the following treatments: control, - Al, and two doses of Al (25 µM and 100 µM AlCl3). The seedlings remained in these conditions for 3 days before sampling for further analyses.

A second experiment, using plants under later developmental stage, under the same conditions, was also conducted. Briefly, 10-day-old seedlings were transferred to the nutrient solution described above. After 18 days of acclimation, the plants were subjected to the treatments: control, -Al, and two doses of Al (25 µM and 100 µM AlCl3) for 7 days, followed by sampling for further analyses. For biochemical and molecular analyses, shoot and root samples were collected at midday, flash-frozen in liquid nitrogen (N_2_) and stored at −80 °C until further analysis. Root length was measured from the hypocotyl to the tip of the seminal root using a scanner (Hewlett Packard Scanjet G2410, Palo Alto, California, USA). The resulting images were processed in ImageJ to calculate root growth rates. Thus, the relative growth was calculated by dividing growth at Al stress per growth at control conditions.

### 2.2 Histochemical Al localization

Root tips were immersed in a 0.2% (w/v) iron hematoxylin solution, containing sodium iodide (NaIO3) for 15 minutes (Souza et al., 2016). Subsequently, the roots were washed in aerated deionized water for 15 minutes to remove excess staining. Samples were observed and photographed under a stereomicroscope (Zeiss model Stemi 2000-C).

### 2.3 Anatomical assays and immunolocalization of Al

Root tip samples (∼1 cm) were collected and fixed in 50% FAA (formaldehyde 37%, glacial acetic acid, and 50% ethanol, 1:1:18 v/v/v) for 72 hours, then stored in 70% ethanol. The samples were dehydrated through a graded ethanol series and embedded in methacrylate resin (Historesin, Leica Instruments, Heidelberg, Germany). Transverse and longitudinal sections (5 µm) were prepared using an automatic rotary microtome (RM2155, Leica Microsystems Inc., Deerfield, USA). Sections were stained with toluidine blue for general anatomical observation. For aluminium detection, sections were stained with 0.5% Chrome Azurol S (Kukachka and Miller, 1980). After 1 and 60 minutes of incubation in these reagents, respectively, sections were washed in distilled water. Permanent slides were prepared using Permount and observed under a photomicroscope (model AX70 TRF, Olympus Optical, Tokyo, Japan) equipped with an AxioCam Zeiss camera.

### 2.4 Histochemical ROS determination

Shoot and root apex samples were collected to qualitatively assess hydrogen peroxide (H2O2) and superoxide (O^2-^) levels using histochemical staining, as described by Kong *et al*. (2011), with modifications to exposure time and reagent concentration. Briefly, Nitroblue Tetrazolium (NBT) at 0.1 mg ml^-1^ was used to detect O^2-^, while 3,3’-diaminobenzidine (DAB) at 1.0 mg ml^-1^ was used to detect H2O2. Samples were exposed to NBT and DAB solutions for 1 and 3 hours, respectively. After staining, the solutions were removed, and a fixation solution (ethanol: acetic acid: glycerol 3:1:1) was added to cover all samples. Samples were stored until analysis and photographed under a stereomicroscope (Zeiss Stemi 2000-C model).

### 2.5 Measurements of gas exchange and chlorophyll fluorescence

Gas exchange parameters were measured simultaneously with chlorophyll *a* (Chl *a*) fluorescence, as described by Nascimento et al. (2021). Analyses were performed using an open-flow infrared gas exchange analyser system (LI-6400XT; LI-COR Inc., Lincoln, NE, USA) equipped with an integrated fluorescence chamber (LI-6400-40; LI-COR Inc.). Instantaneous gas exchange was measured after 1 h illumination during the light period under 1000 µmol photons m^−2^ s^−1^, corresponding to the photosynthetically active photon flux density (PPFD) saturation level for leaves as determined by Nascimento et al. (2021). The reference CO2 concentration was set at 400 µmol CO2 mol^−1^ air. Measurements were performed using a 2 cm^2^ leaf chamber at 25 °C, with a 0.5 stomatal ratio (amphistomatic leaves). The leaf-to-air vapor pressure deficit was kept at 1.2 kPa, and 10% of the PPFD was provided as blue light to optimize stomatal aperture.

Initial fluorescence emission (*F*0) was obtained by illuminating dark-adapted leaves (2 h) with a weak modulated measuring beam (0.03 µmol photons m^−2^ s^−1^). A saturating white light pulse (8000 µmol photons m^−2^ s^−1^) was applied for 0.8 s to measure maximum fluorescence (*F*ₘ). The variable-to-maximum chlorophyll fluorescence ratio (*F*ᵥ/*F*ₘ) was calculated as *F*v/*F*m = [(*F*m–*F*0)/*F*m)]. The efficiency of excitation energy capture by open photosystem II (PSII) reaction centre (*F*v’/*F*m’) was estimated following Logan *et al*. (2007), whereas the actual PSII photochemical efficiency (φPSII) was estimated as φPSII=(*F*m’-*F*s)/*F*m’ according to Genty *et al*. (1989). Dark respiration (R*d*) was measured after 2 h in the dark (night period) using the same gas exchange system described above and was divided by two (R*d*/2) to estimate the mitochondrial respiration rate during the light (R*d*).

### 2.6 Biochemical analyses

Metabolites were extracted by grinding tissue in liquid nitrogen followed by the immediate addition of methanol. Photosynthetic pigments were determined as described by Porra et al. (1989). Starch, sucrose, fructose, and glucose levels in the leaf tissue were determined following Cross et al. (2006). Malate and fumarate were measured as described by Nunes-Nesi et al. (2007). Total protein content was determined using the Bradford (1976) method, while total amino acid levels were quantified following Cross et al. (2006). Metabolite profiling was carried out as described by Lisec et al. (2006), with modifications. Briefly, approximately 25 mg of homogenized leaf tissue was aliquoted into tubes and extracted in 100% methanol with an internal standard (0.2 mg ribitol mL^− 1^ water). The derivatization and sample injection steps were performed exactly as described before (Lisec et al., 2006). Peak detection, retention time alignment, and library matching were carried out using the Target Search R-package. Metabolites were identified by comparison with database entries of authentic standards (Kopka et al., 2005). Metabolite identification and annotation followed the reporting recommendations described in Fernie et al. (2011).

## 3. RESULTS

### 3.1 Vegetative growth is differentially affected by Al toxicity in auxin perception mutants

To determine whether auxin signalling influences growth responses under Al stress conditions, we used *Solanum lycopersicum* cv. Micro-Tom (MT) mutants with enhanced (*entire*) and reduced (*diageotropica, dgt)* auxin signalling. These plants were subjected to two Al doses (25 and 100 µM) for three days. Growth was assessed during early development (15 days after germination, DAG) and late development (35 DAG) to examine whether plant stage combined with auxin perception affects Al tolerance. During the early development, root growth of the *entire* mutant as greater than that of the WT under the highest Al dose, namely 100 µM Al (Figure 1A-B). By exposing plants to 100 µM Al at late development, root growth in WT and *dgt* plants was reduced by 53%, while the *entire* mutant exhibited less inhibition, with root growth reduced by only 18% (Figure 2A-C). Altogether, these results suggest that the *entire* mutant exhibits higher Al tolerance. This prompted us to investigate whether the plants displayed differential Al distribution across root sections.

**Figure 1.**
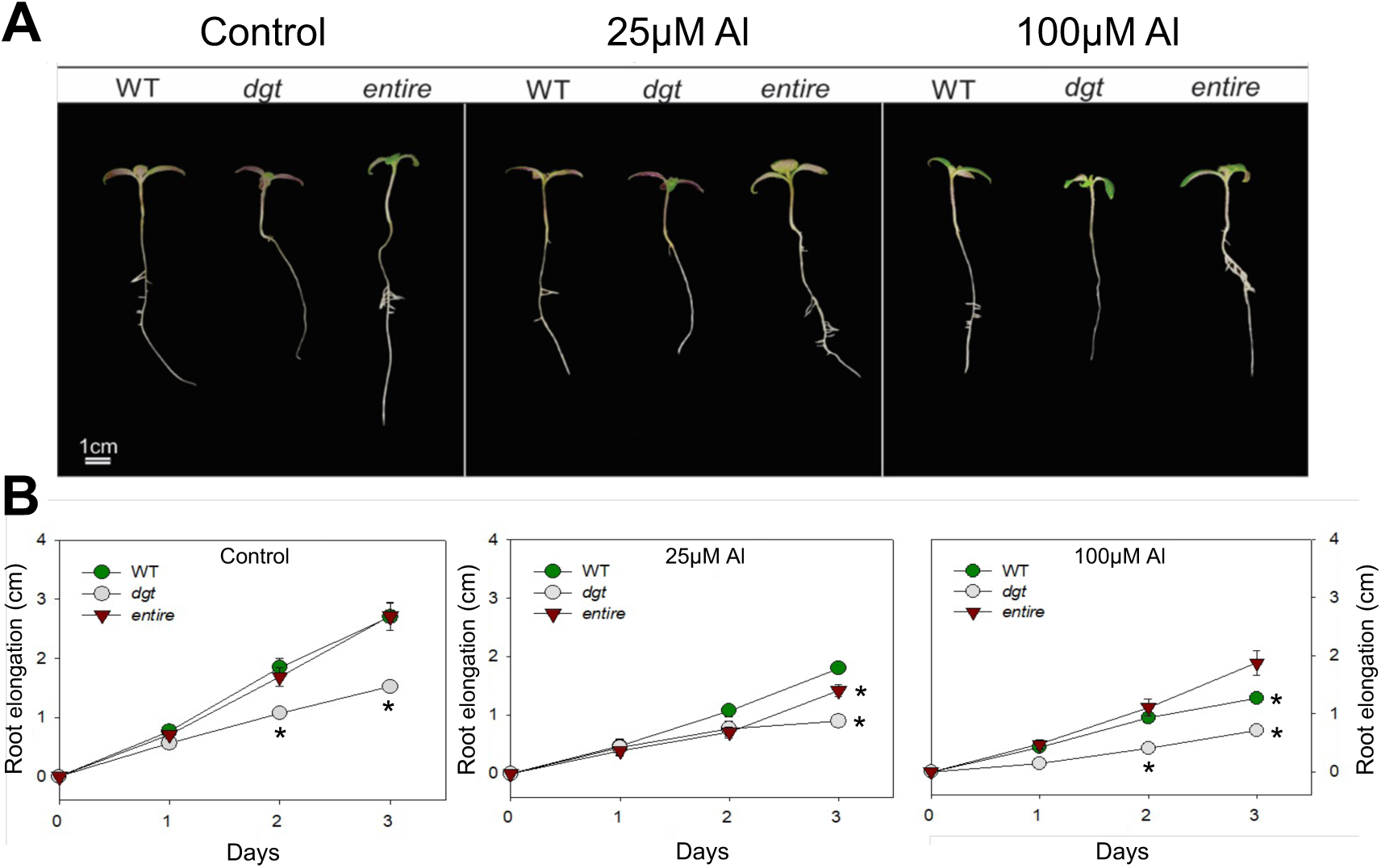
Increased auxin signalling in *entire* mutant enhances Aluminium (Al) tolerance in *Solanum lycopersicum*. Seeds of the wild-type (WT), *diageotropica* (*dgt*), and *entire* mutant lines, all derived from the cultivar Micro-Tom, were germinated and grown for 15 days after germination. Subsequently, these seedlings were subjected to 0 μM (control), 25 μM, and 100 μM Al for three days. **(A)** Phenotypes of plants after three days of exposure to Al stress. **(B)** Root elongation during the period of exposure to Al stress. The experimental conditions were performed as detailed in the Materials and Methods section. Data represent eight individual plants. Values are expressed as means ± SE (n = 8) and were compared using the two-sided Student’s *t*-test. Differences were considered statistically significant at *P* < 0.05. Asterisks indicate significant differences compared to the WT at each time point in presence of Al.

**Figure 2.**
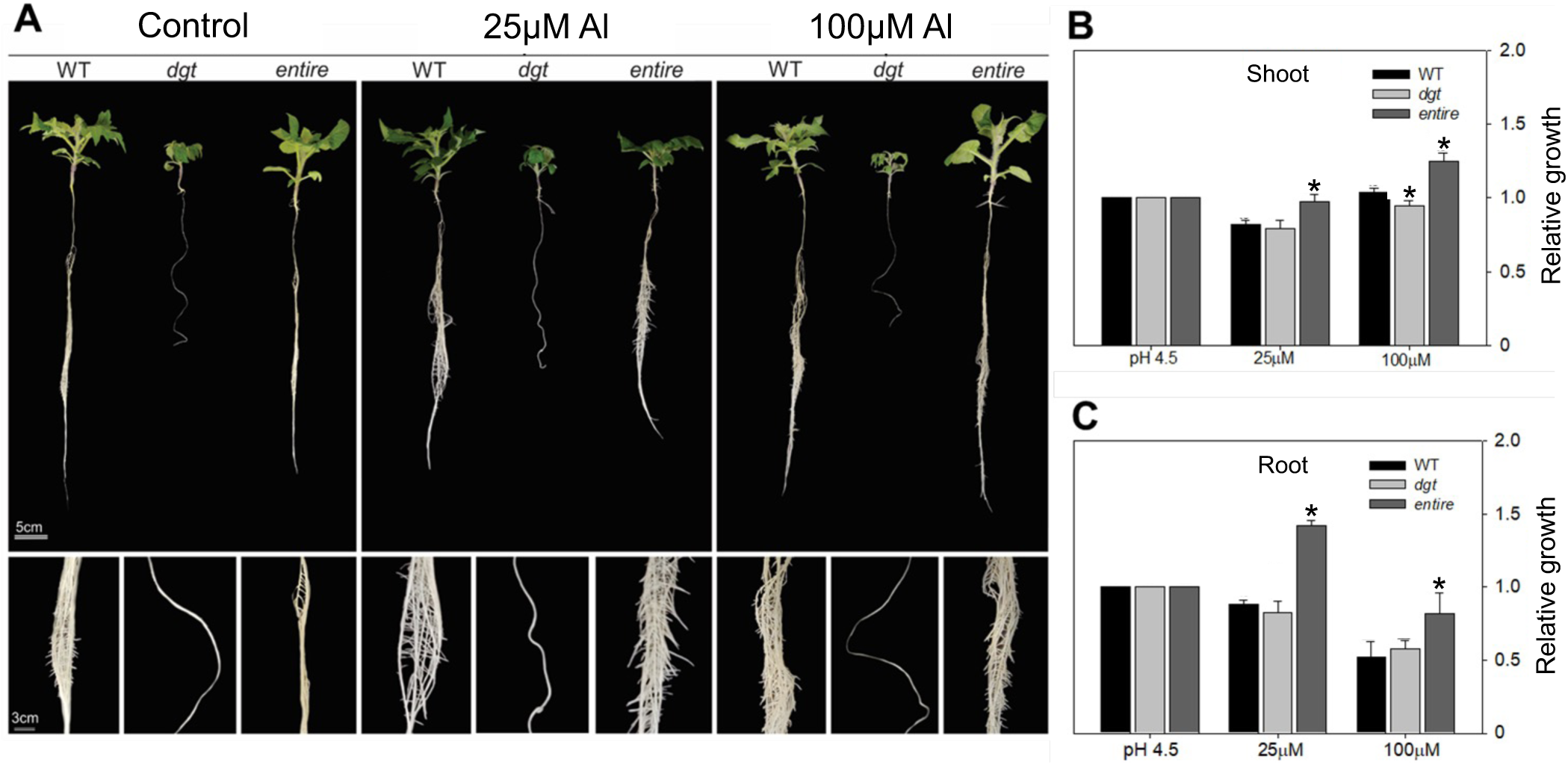
The *entire* mutant exhibits prolonged Aluminium (Al) tolerance. Plants of the wild-type (WT), *diageotropica* (*dgt*), and *entire* mutant lines, all derived from the cultivar Micro-Tom, were germinated and grown for 35 days after germination. Subsequently, these plants were subjected to 0 μM (control), 25 μM, and 100 μM Al for three days. **(A)** Phenotypes of plants after three days of exposure to Al stress. **(B-C)** Relative shoot and root growth, showing data normalized to the average response of each genotype under control conditions (pH 4.5). Variations in growth under Al treatment reflect the impacts of Al stress on shoot (B) and root (C) growth for each genotype. Growth conditions were conducted as described in the Materials and Methods section. Data represent eight individual plants. Values are expressed as means ± SE (n = 8) and were compared using the two-sided Student’s *t*-test. Differences were considered statistically significant at *P* < 0.05. Asterisks indicate significant differences compared to the WT at each Al concentration.

### 3.2 Al immunolocalization on roots of tomato lines

The immunolocalization of Al in roots was evaluated using hematoxylin staining, which revealed staining only in plants treated with Al (Figure 3). Reduced auxin perception associated with the *dgt* mutant resulted in intense root staining, particularly in root tip regions, even at a low Al dose of 25 µM (Figure 3). In contrast, the *entire* mutant exhibited significantly less staining, with Al accumulation observed on the root surface only at the higher Al dose of 100 µM (Figure 3). These results suggest that higher auxin perception in the *entire* mutant at low Al concentrations reduces Al accumulation on root surfaces.

**Figure 3.**
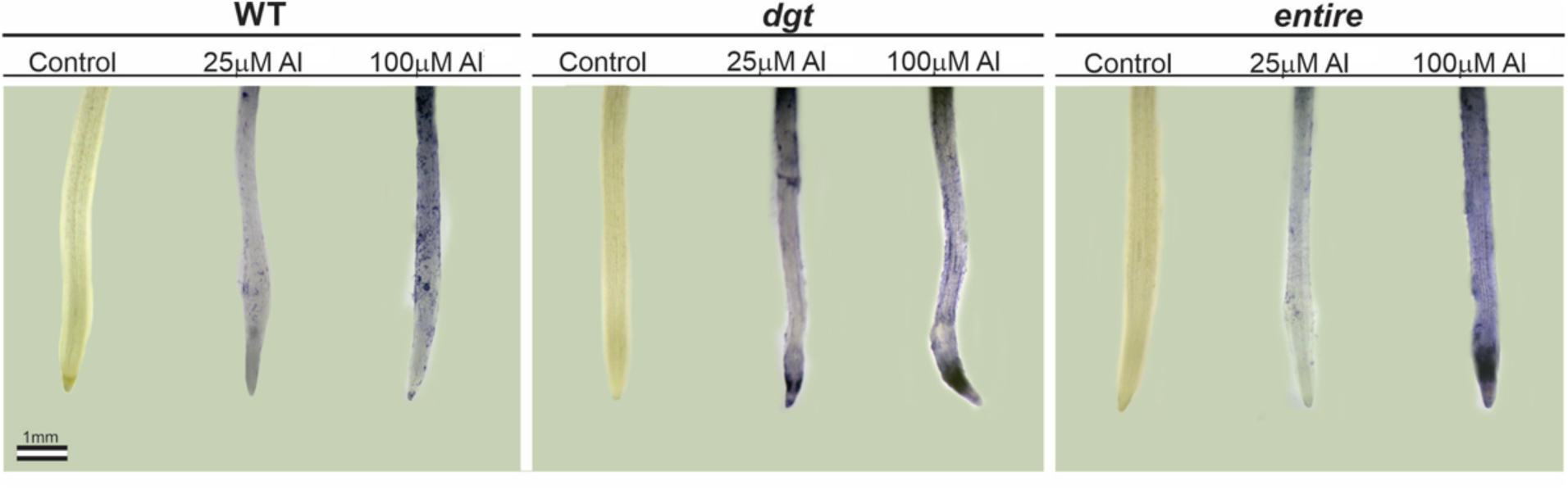
Hematoxylin staining of three *Solanum lycopersicum* lines exposed to Al stress. Seeds of wild-type (WT), *diageotropica* (*dgt*), and *entire* mutant lines, all derived from the cultivar Micro-Tom, were germinated, and plants were grown for 35 days after germination. Subsequently, these seedlings were subjected to 0 μM (control), 25 μM, and 100 μM Al for three days. The images show root apices 18 days after germination, where Aluminium (Al) is stained purple. **Scale bar = 1 mm.**

### 3.3 Changes in ROS staining for tomato lines under aluminium stress

Histochemical assays using Nitro Blue Tetrazolium (NBT) and Diaminobenzidine (DAB) provided a qualitative assessment of superoxide (O2^-^) and hydrogen peroxide (H2O2), respectively. These assays were performed in the shoots and roots of tomato genotypes (35 DAG) with varying auxin sensitivities. Under control conditions and at 25 µM Al, weak staining was observed in the aerial parts of all genotypes for both NBT and DAB assays (Figure 4A-B). However, exposure to 100 µM Al resulted in intense staining in all genotypes, particularly in WT and *entire* plants, which showed even stronger coloration for DAB, but the differences were not clear for NBT (Figure 4A-B). Notably, the *dgt* mutant exhibited less intense staining, indicating a lack of redox response to Al stress, which may have contributed to its absence of response and higher sensitivity to Al stress.

**Figure 4.**
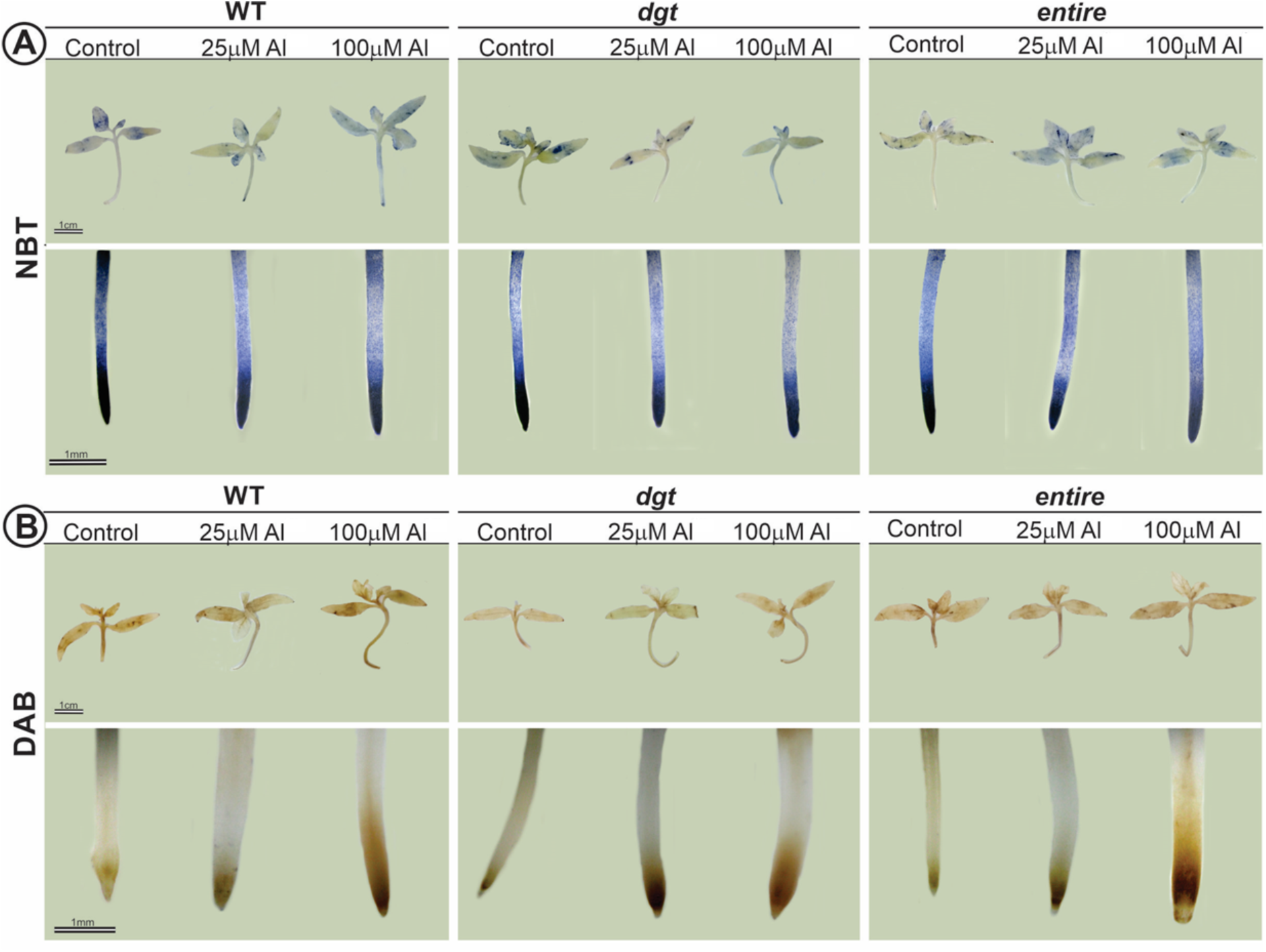
Histochemical assay for the immunolocalisation of reactive oxygen species (ROS) in three *Solanum lycopersicum* lines exposed to Al stress. Seeds of wild-type (WT), *diageotropica* (*dgt*), and *entire* mutant lines, all derived from the cultivar Micro-Tom, were germinated, and plants were grown for 35 days after germination. Subsequently, these seedlings were subjected to 0 μM (control), 25 μM, and 100 μM Aluminium (Al) for three days. Plants exposed to control and Al stress conditions for three days were stained with Nitro Blue Tetrazolium (NBT) to detect O₂ (**A**) and with Diaminobenzidine (DAB) to localise H₂O₂ (**B**). Treated plants were compared with their respective controls at the time of harvesting.

### 3.4 Gas exchanges and metabolism

To understand whether auxin signalling influences leaf physiology in tomato plants under Al stress, we characterized gas exchange and chlorophyll *a* fluorescence in plants at late development (35 DAG). The *dgt* mutant showed the lowest values for CO2 assimilation net (*AN*), stomatal conductance (*gs*), and transpiration (*E*), even in the absence of Al stress. These differences were generally maintained under Al toxic conditions (Table I). Dark respiration (*Rd*) was also lower in the leaves of the *dgt* genotype under Al stress (Table I). Notably, internal CO2 (Ci) was increased for *entire* and reduced at *dgt* both compared WT plants (Table I). Ci also showed a distinct pattern, with an increase for entire mutant and a decrease for *dgt*. While the *dgt* mutant exhibited the lowest values for most gas exchange parameters, the *entire* mutant remained practically unaffected by Al stress.

**Table I.** Gas exchange and chlorophyll fluorescence *a* of plants exposed to toxic levels of Aluminium (Al). Seeds of wild-type (WT), *diageotropica* (*dgt*), and *entire* mutant lines, all derived from the cultivar Micro-Tom, were germinated, and plants were grown for 35 days after germination. Subsequently, these seedlings were subjected to 0 μM (control), 25 μM, and 100 μM Al for three days. Values are expressed as means ± SE (n = 5) and were compared using the two-sided Student’s *t*-test. Differences were considered statistically significant at *P* < 0.05. Asterisk indicates significant difference compared to the WT at each condition tested.

| Parameters | Treatments |  |  |
| --- | --- | --- | --- |
| | Control | 25$\mu\text{M}$ Al | 100$\mu\text{M}$ Al |
| **$A_N$ (mmol $\text{CO}_2 \text{ m}^{-2} \text{ s}^{-1}$)** |  |  |  |
| wt | 5.59 ± 0.44 | 5.31 ± 0.39 | 5.13 ± 0.89 |
| *dgt* | **3.50 ± 0.41\*** | **3.85 ± 0.35\*** | **4.00 ± 0.81\*** |
| *entire* | 5.41 ± 0.22 | 5.18 ± 0.27 | 5.71 ± 0.34 |
| **$g_s$ (mol $\text{H}_2\text{O m}^{-2} \text{ s}^{-1}$)** |  |  |  |
| wt | 0.24 ± 0.03 | 0.30 ± 0.02 | 0.28 ± 0.10 |
| *dgt* | **0.10 ± 0.01\*** | **0.15 ± 0.01\*** | **0.14 ± 0.07\*** |
| *entire* | **0.27 ± 0.01\*** | 0.33 ± 0.02 | 0.28 ± 0.03 |
| **$E$ (mmol $\text{H}_2\text{O m}^{-2} \text{ s}^{-1}$)** |  |  |  |
| wt | 2.08 ± 0.23 | 2.52 ± 0.34 | 2.61 ± 0.70 |
| *dgt* | 1.09 ± 0.20 | 1.47 ± 0.09 | 1.50 ± 0.52 |
| *entire* | 2.34 ± 0.19 | 2.71 ± 0.31 | 2.81 ± 0.15 |
| **$WUE_i$ ($A_N/g_s$)** |  |  |  |
| wt | 24.14 ± 1.61 | 18.16 ± 2.02 | 23.31 ± 50.99 |
| *dgt* | **35.68 ± 2.57\*** | **25.77 ± 3.16\*** | **39.46 ± 114.81\*** |
| *entire* | **20.06 ± 0.41\*** | 15.91 ± 0.61 | 20.51 ± 125.47 |
| **$R_d$ (mmol $\text{CO}_2 \text{ m}^{-2} \text{ s}^{-1}$)** |  |  |  |
| wt | 2.57 ± 0.15 | 3.25 ± 0.35 | 3.10 ± 0.25 |
| *dgt* | **1.97 ± 0.11\*** | **2.65 ± 0.04\*** | 3.07 ± 0.29 |
| *entire* | **3.10 ± 0.39\*** | 4.09 ± 0.28 | 3.49 ± 0.35 |
| **$ETR$** |  |  |  |
| wt | 44.87 ± 0.39 | 43.93 ± 0.44 | 42.56 ± 1.40 |
| *dgt* | 41.49 ± 1.14 | 41.65 ± 0.68 | 40.08 ± 1.17 |
| *entire* | 44.44 ± 0.26 | 43.88 ± 0.53 | 43.30 ± 0.43 |
| **$F_v/F_m$** |  |  |  |
| wt | 0.78 ± 0.00 | 0.78 ± 0.00 | 0.78 ± 0.00 |
| *dgt* | 0.76 ± 0.00 | 0.76 ± 0.00 | 0.78 ± 0.00 |
| *entire* | 0.77 ± 0.00 | 0.78 ± 0.00 | 0.78 ± 0.00 |
| **$C_i$** |  |  |  |
| wt | 350.64 ± 2.51 | 360.61 ± 3.62 | 345.80 ± 9.59 |
| *dgt* | **333.32 ± 3.57\*** | **349.36 ± 5.63\*** | 326.02 ± 18.55 |
| *entire* | **357.50 ± 0.34\*** | 364.28 ± 1.40 | **355.50 ± 2.10\*** |

Abbreviations: $\text{CO}_2$ assimilation net ( $A_N$ ), stomatal conductance ( $g_s$ ), transpiration ( $E$ ), intrinsic water-use efficiency ( $WUE_i$ ), dark respiration ( $R_d$ ), electron transport rate ( $ETR$ ), maximum quantum yield of photosystem II ( $F_v/F_m$ ), and internal $\text{CO}_2$ ( $C_i$ ).

To investigate the link between auxin perception and plant metabolism under Al stress, a detailed metabolic characterization was performed. Al stress favoured the accumulation of glucose, fructose, and starch in the shoots of the *dgt* mutant, while its roots exhibited higher levels of sucrose and amino acids compared to WT plants (Figures 5-6). The organic acid malate, which plays a fundamental role in Al tolerance, accumulated in both shoots and roots of *dgt* plants. However, at 100 µM Al, malate levels were significantly reduced in the shoots of *dgt* plants (Figure 6B). Meanwhile, under this same Al concentration, the *entire* mutant exhibited a remarkable reduction in malate levels, which may contribute to its Al tolerance (Figure 6B).

**Figure 5.**
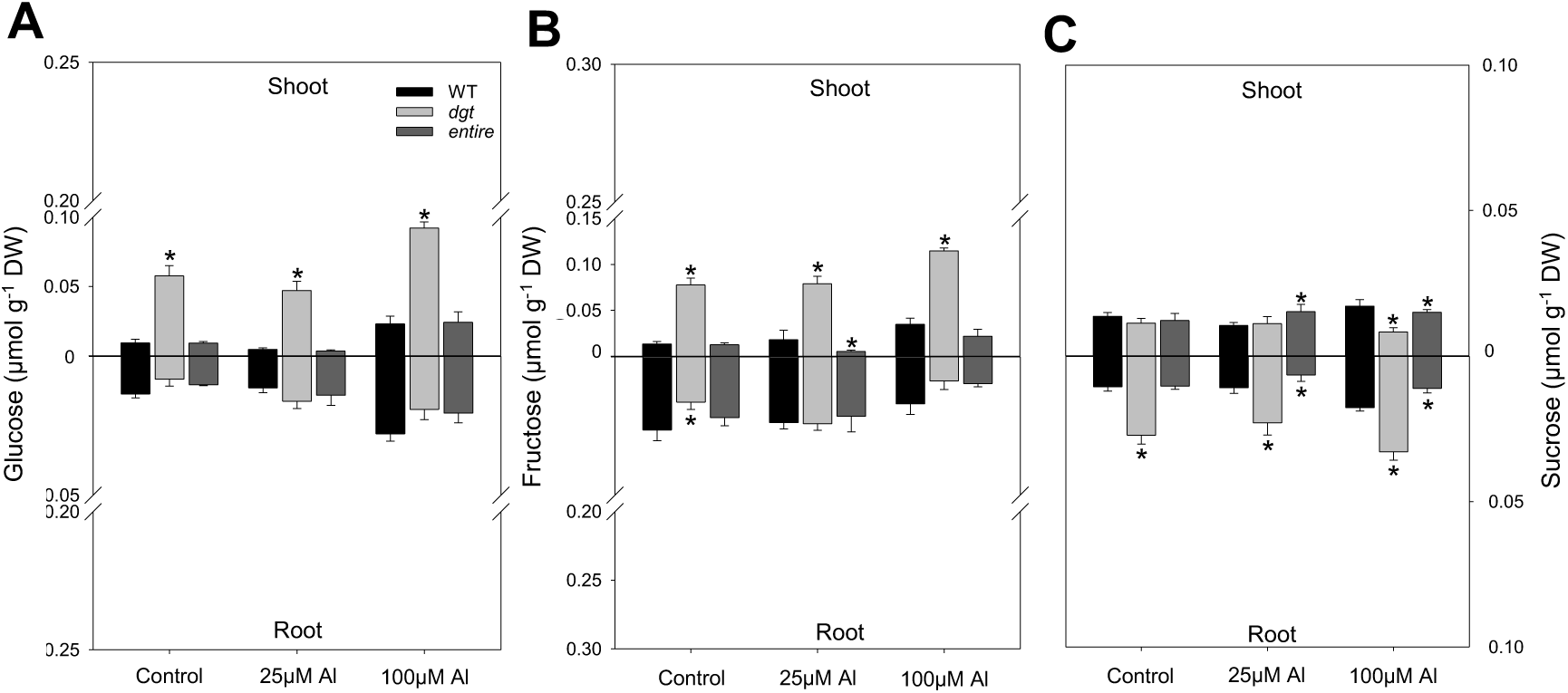
Sugar levels in shoots and roots of tomato plants exposed to Aluminium (Al) toxicity. The levels of **(A)** glucose, **(B)** fructose, and **(C)** sucrose were determined in plants 35 days after germination, following exposure to 0 μM (control), 25 μM, and 100 μM Al for three days. Values are expressed as means ± SE (n = 5) of individual plants and were compared using the two-sided Student’s *t*-test. Differences were considered statistically significant at *P* < 0.05. Asterisks indicate significant differences compared to the WT under each condition tested.

**Figure 6.**
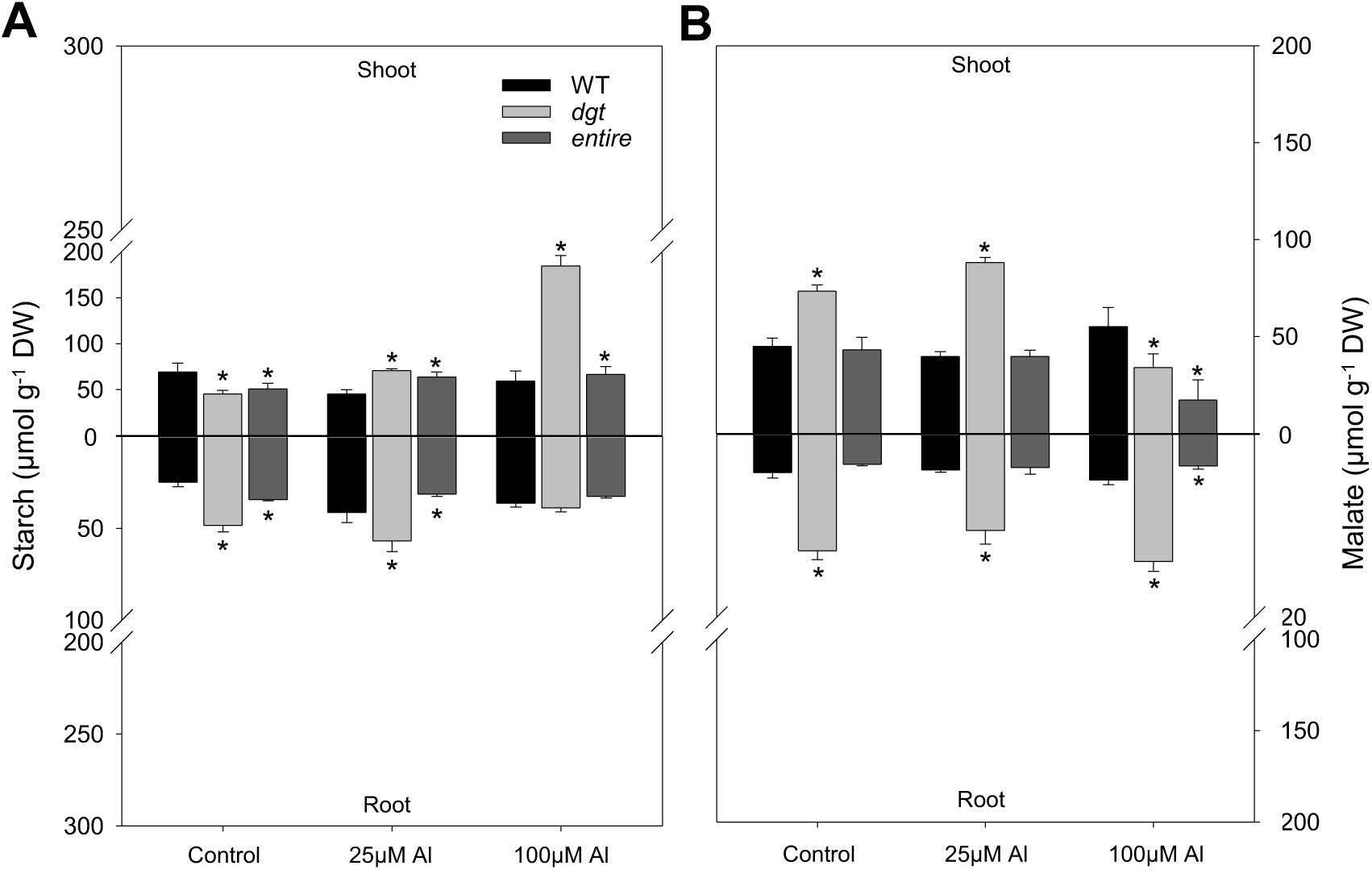
Starch and malate levels in shoots and roots of tomato plants exposed to Aluminium (Al) toxicity. The levels of **(A)** starch and **(B)** malate were determined in plants 35 days after germination, following exposure to 0 μM (control), 25 μM, and 100 μM Al for three days. Data represent five individual plants. Values are expressed as means ± SE (n = 5) and were compared using the two-sided Student’s *t*-test. Differences were considered statistically significant at *P* < 0.05. Asterisks indicate significant differences compared to the WT under each condition tested.

Since the major impacts of Al toxicity on growth and malate occurred in the roots, a metabolite profile was performed to characterize root metabolism in the genotypes used. This analysis revealed 45 successfully annotated compounds, including amino acids, organic acids, sugars, and metabolic intermediates (Figure 7 and Table S1). Changes in compounds associated with carbon and nitrogen metabolism were observed due to differences in auxin signalling and exposure to Al stress (Figure 7, Table S1). In the absence of Al stress, minor metabolic differences were observed between WT and *entire* plants. However, in *dgt* plants, 10 individual amino acids, including alanine, arginine, aspartate, β-alanine, phenylalanine, GABA, glutamine, homoserine, lysine, methionine, threonine, and valine, were significantly increased (Figure 7, Table S1). Similarly, organic acids such as aconitate, glycerate, gluconate, malate, nicotinate, saccharopine, and succinate, as well as sugars like fructose, glucose, and sucrose, and sugar alcohols such as glycerol, mannitol, mannose, and myo-inositol, were higher in *dgt* plants compared to the other genotypes (Figure 7, Table S1).

**Figure 7.**
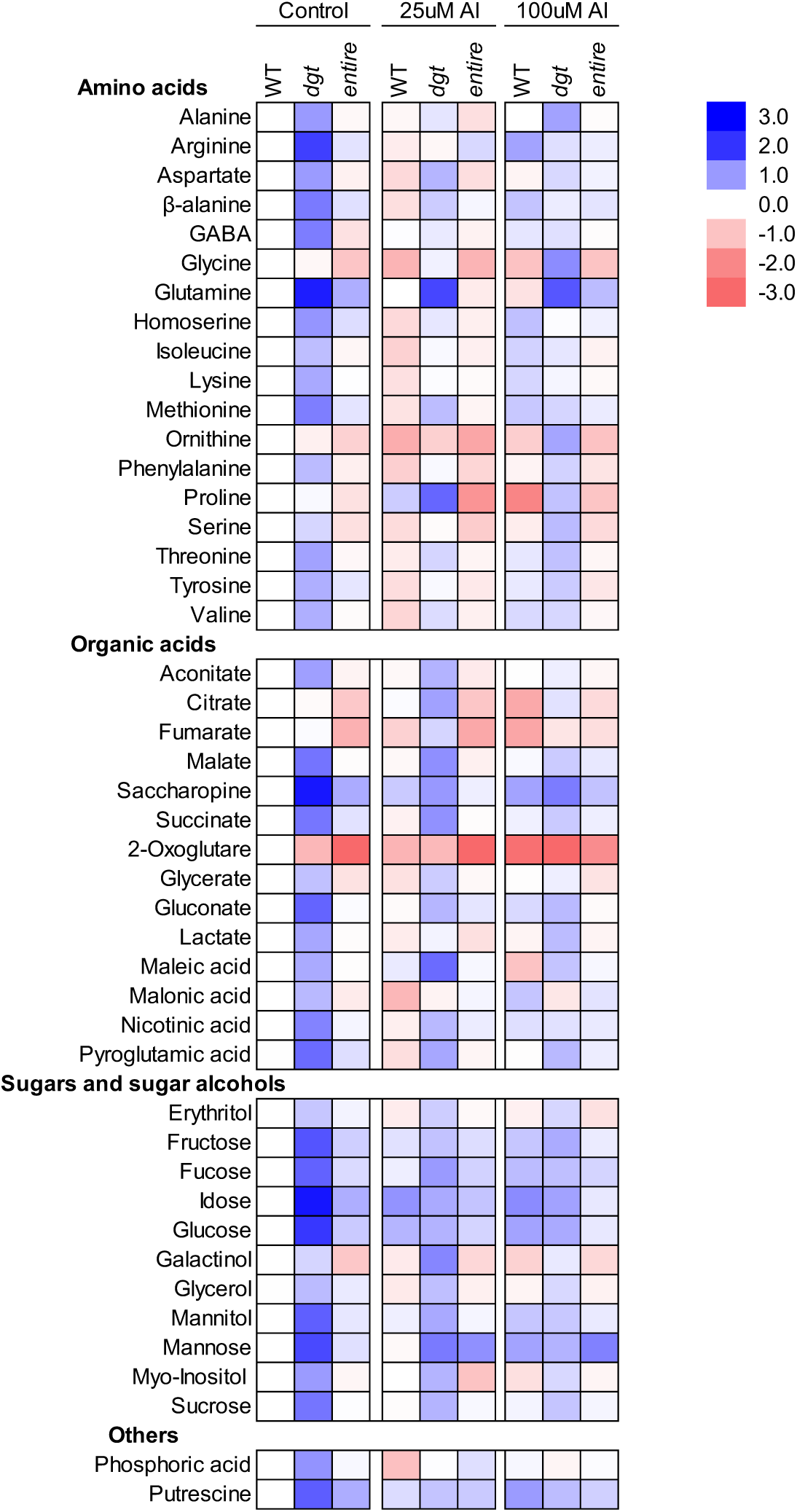
Heatmap representing changes in root metabolite levels in *Solanum lycopersicum* lines exposed to Aluminium (Al) stress. Metabolite levels were normalized to wild-type (WT) control plants grown under control (0 μM Al) conditions. Values were According to the lateral scale, red colour intensity indicates a reduction in metabolite levels, while blue colour intensity indicates an increase. Metabolites were determined as described in “Materials and Methods.” Values presented are means of six biological replicates. Further statistical details can be found in Supplementary Table S1.

When exposed to Al stress, *dgt* plants exhibited significant reductions in several metabolites. Seven amino acids, including arginine, aspartate, β-alanine, phenylalanine, homoserine, lysine, and methionine, were reduced under Al stress (Figure 7, Table S1). Similarly, levels of maleate, aconitate, gluconate, lactate, malate, nicotinate, saccharopine, and succinate were lower in *dgt* plants under Al stress (Figure 7, Table S1). Sugars such as fructose, glucose, sucrose, and sugar alcohols (e.g., mannitol, myo-inositol) also showed notable reductions in *dgt* plants, particularly at 100 µM Al (Figure 7, Table S1). Interestingly, proline levels tended to increase in *dgt* plants in the presence of 25 µM Al compared to other genotypes (Figure 7). Additionally, *dgt* plants exposed to 100 µM Al showed increases in serine and threonine levels (Figure 7).

### 3.5 Root microanatomy

By using microscopy of the root apical meristem (RAM) and Chrome Azurol S staining to immunolocalize Al, we observed significant changes in root structure due to Al treatment. In the root differentiation zone of WT and *dgt* plants, larger vacuoles were observed, correlating with increased cell differentiation (Figure 8A-C), a characteristic response of Al-sensitive genotypes. In contrast, *entire* plants exhibited a reduced number of differentiated cells in the root meristematic zone, with these cells showing fewer evident nuclei (Figure 8). Al was preferentially immunolocalized in the nucleus, with more intense staining in WT and *dgt* genotypes compared to *entire*, through the reduced number of blue staining dots at *entire* root tips (Figure 8). Therefore, these findings support the notion of a higher Al tolerance for the *entire* mutant.

**Figure 8.**
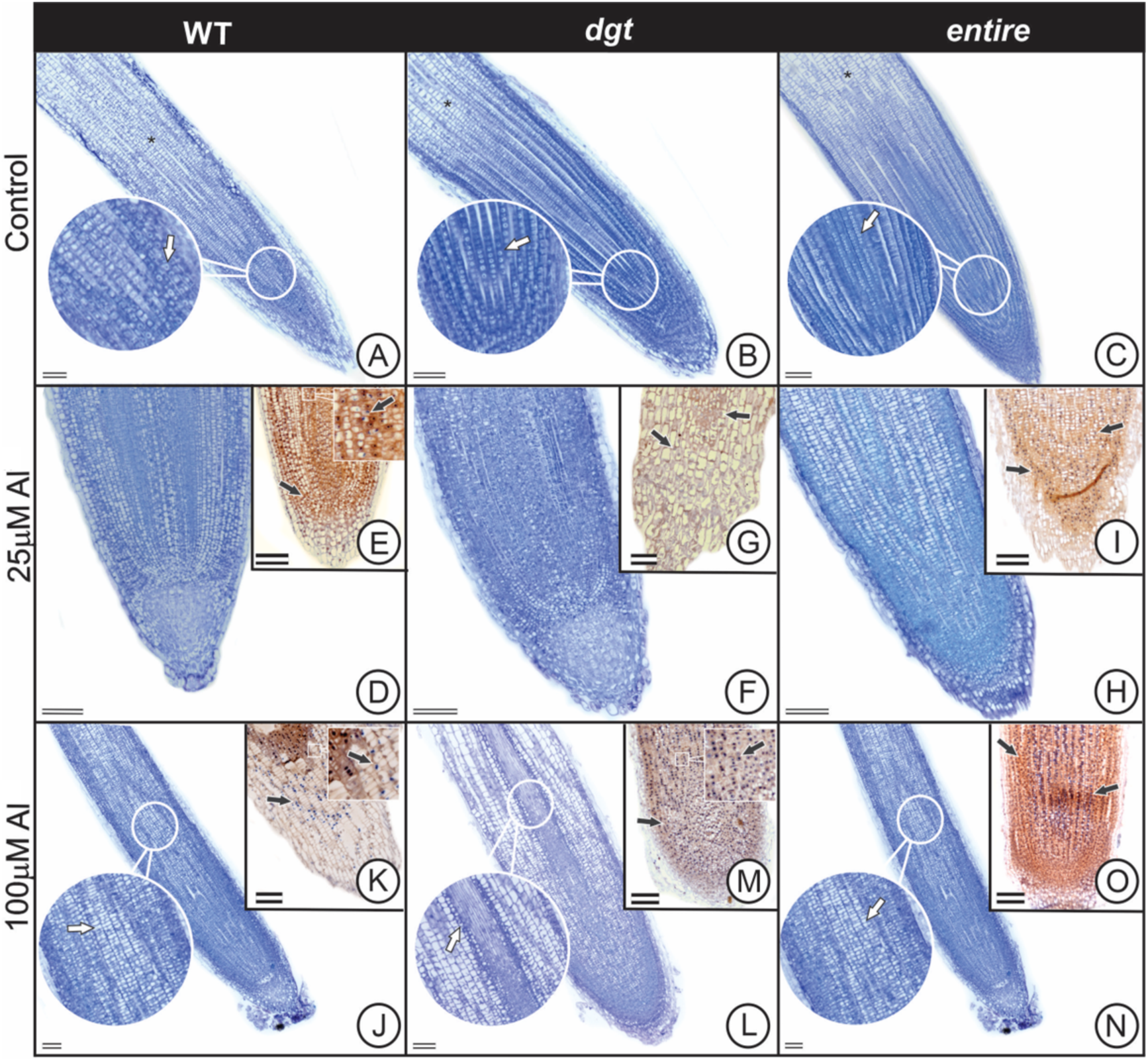
Al stress induces changes in the cellular structures of root apices in tomato plants with differential auxin signalling. Representative images of root apex anatomy in adult plants (35 days after germination) with reduced (*diageotropica*, *dgt*) or increased (*entire*) auxin signalling, under different concentrations of AlCl₃ (control, 25 and 100 μM), compared with wild-type (WT) plants (*Solanum lycopersicum* cv. Micro-Tom). Images shown in **E, G, I, K, M, O** refers to the immunolocalization of Aluminium (Al) in root apices using Chrome Azurol S stain. The blue staining, indicated by black arrows, represents a positive reaction to Chrome Azurol S. **Scale bar = 1 mm.**

## 4. DISCUSSION

An increasing concern for humanity is the development of crops with enhanced nutrient use efficiency that can grow in acidic soils (Siqueira et al., 2023). Al toxicity has caused agricultural problems for centuries, reducing food production by limiting the use of croplands in many developing regions. Historically, agricultural expansion was limited in areas with elevated levels of Al in the soil. For instance, the Central-western region of Brazil faced this phenomenon until the 1980s when advances in soil pH correction techniques enabled agricultural expansion in this region. The major mechanisms enabling plant growth under Al toxicity include the exudation of OAs from root cells into the rhizosphere and alterations in DNA checkpoints, both of which promote root cell division under Al stress (Siqueira et al., 2018; Siqueira et al., 2022a). Since these mechanisms are fundamental for Al tolerance and individual genetic manipulations of these pathways have proven insufficient to overcome Al stress, additional mechanisms are likely involved in regulating Al tolerance (Nunes-Nesi et al., 2014; Siqueira et al., 2022a; Pereira-Lima et al., 2023). Some genotypes under nutritional disturbances exhibit parallel imbalances in hormone homeostasis and primary metabolism (Song and Liu, 2015). Auxin is a promising candidate for orchestrating metabolism and cell division during Al stress conditions (Kopittke, 2016; Zhang et al., 2018; Wang et al., 2016a). To elucidate the interaction between auxin signalling and Al tolerance, we characterized tomato responses to Al toxicity using mutants with either increased (*entire*) or reduced (*dgt*) auxin signalling. While both genes are relatively well-characterized in modulating growth and metabolism of tomatoes (Kelly and Bradford, 1986; Coenen et al., 2002; Balbi and Lomax, 2003; Wang et al., 2005; Goldental-Cohen et al., 2017; Batista-Silva et al., 2019), our work demonstrates an intrinsic relationship between auxin signalling and Al tolerance. Regardless of the tomato genotype, Al stress reduced root elongation and induced specific alterations in root anatomy. Importantly, differential auxin signalling contributed to contrasting responses to Al toxicity. Specifically, amplified auxin perception in the *entire* mutant increased Al tolerance, whereas deficient auxin signalling in *dgt* exacerbated the negative effects of Al toxicity.

Auxin signalling modulates plant growth by regulating pathways related to development and metabolism, including cell proliferation and division in shoots and roots (Batista-Silva et al., 2018; Ivanova et al., 2014; Poomam et al., 2015). The *dgt* and *entire* mutants exhibit clear growth alterations compared to WT plants, highlighting reduced apical dominance in both mutants and fewer lateral ramifications in *dgt* plants (Oh et al., 2006; Ivanchenko et al., 2015). The maintenance of growth under Al toxicity does not appear to be related to apical dominance but rather to lateral branching, as *dgt* lacks lateral ramifications and is Al sensitive, while *entire* retains root branches, allowing it to tolerate Al stress (Fig. 1-3). The root transition zone is the primary site for Al stress perception, and phytohormones, including auxin, mediate Al-induced inhibition of root growth (Yang et al., 2014; Liu et al., 2016; Yang et al., 2017). Al can affect polar auxin transport and intracellular auxin distribution, leading to its accumulation in the root transition zone and ultimately inhibiting root growth (Li et al., 2021; Siqueira et al., 2022c). The differences in Al tolerance among WT, *dgt*, and *entire* most likely derives from auxin sensitivity and perception. Thus, the reduced auxin sensitivity and perception in *dgt* culminates in a limited Al tolerance, whereas higher auxin sensitivity and perception in the *entire* mutant enhance its Al tolerance.

Differential auxin perception modifies various aspects of vegetative growth in tomatoes, such as the reduced stature and leaf area observed in *dgt* and the conversion from compound to simple leaves in *entire* (Zobel, 1974; Wang et al., 2005; Batista-Silva et al., 2018, 2019). These traits explain the small fruits with reduced biomass in *dgt* plants (Batista-Silva et al., 2022) and highlight the importance of understanding the major factors impacted by Al toxicity. Consistent with Batista-Silva et al. (2019), our results showed that *dgt* mutant plants had lower *AN*, *gs*, and *E* even in the absence of Al stress, with these differences persisting under Al toxicity (Table I). During Al stress, respiratory rates in leaves and roots can distinguish sensitive from Al-tolerant genotypes, with Al tolerance often associated with the maintenance of respiration (Siqueira et al., 2020; Pereira-Lima et al., 2023). Our findings support this hypothesis, as *entire* mutant consistently exhibited the highest respiration rates in leaves across all treatments (Table I), contributing to its elevated Al tolerance compared to WT and *dgt* plants. Auxin is known to promote ATP utilization by regulating cell expansion and division (French and Beevers, 1953; Tivendale and Millar, 2022). Recent studies suggest a direct relationship between auxin and mitochondrial respiration, as mutations in mitochondrial-specific genes can disrupt auxin signalling (Liu et al., 2019; Zhang et al., 2014; Ohbayashi et al., 2019). For example, knockdown of *SDHAF2* (*succinate dehydrogenase assembly factor 2*) increases root sensitivity to acidic pH (5.0), repressing growth due to auxin hyperaccumulation and unbalanced mitochondrial respiration (Tivendale et al., 2021). Thus, the sensitivity of *dgt* mutant plants to Al stress may be related to its reduced respiration, while qualitative metabolic alterations further influence its responses to Al toxicity.

Inefficient growth under adverse conditions often results in carbohydrate accumulation, as efficient use of photoassimilates is critical for abiotic stress tolerance (Rossi et al., 2015; Nunes-Nesi et al., 2016; Fonseca-Pereira et al., 2022). Starch accumulation, for example, can signal metabolic constraints, limiting growth-related reactions (Ruan, 2012). Auxin is known to downregulate starch accumulation, but *dgt* plants degrade less starch at night, leading to significant accumulation of sugars and organic acids (Miyazawa et al., 1999; Batista-Silva et al., 2018). Consistent with prior studies (Siqueira et al., 2020; Pereira-Lima et al., 2023), our results revealed higher levels of sugars and starch in both shoots and roots of *dgt* plants under Al stress, indicating that reduced growth in these plants may partly result from a diminished ability to mobilize carbon for growth. In shoots, the highest malate concentration was observed in WT and *dgt* plants under control and 25µM Al conditions, with *dgt* plants maintaining elevated malate levels across all conditions. In contrast, *entire* and WT plants exhibited smaller metabolic changes, likely due to the maintenance of respiration and the TCA cycle homeostasis (Batista-Silva et al., 2018). Stress conditions are often accompanied by increased proline production, which acts as a metal chelator and maintains ROS concentrations within acceptable ranges, preventing oxidative bursts (Hayat et al., 2012). Anatomical analyses showed that Al toxicity caused disarrangements in nuclei and cytoplasm organization in all genotypes under higher Al concentrations. Remarkably, at 100µM Al, *entire* plants exhibited fewer differentiated cells in the meristematic zone, indicating that cell division was not compromised. Together, these findings suggest that increased auxin perception support elevated Al tolerance in tomatoes by mitigating the inhibitory effect of Al on root growth. Auxin perception depends on photoperiod, and Al tolerance and nutrient responsiveness are modulated through daylength (Frank et al., 2022; Siqueira et al., 2023). Therefore, future studies should explore how different photoperiods simultaneously modulate auxin signalling and Al tolerance, contributing to the development of the next generation of crops with enhanced tolerance to Al toxicity.

## 5. CONCLUSIONS

Collectively, the results presented here demonstrate that changes in auxin signalling modulate critical responses to Al stress. This study provides a better understanding of the mechanisms employed by tomato genotypes with differential auxin signalling to mitigate or avoid Al toxicity, particularly in the root elongation region. However, further studies are needed to elucidate the mechanisms of Al absorption in response to differential auxin signalling, including changes in gene expression under stress. These findings highlight the importance of auxin signalling in regulating anatomical, molecular, and physiological traits important for plant growth and function, especially in crops of agronomic interest. Modulating auxin signalling pathways offers a promising opportunity to improve the performance of crops of agronomic interest cultivated in acidic soils affected by Al stress.

## Acknowledgments

This work was supported by the National Council for Scientific and Technological Development (CNPq-Brazil) [Grant 151020/2024–8; 407276/2021–1 and 406455/2022–8], and the Foundation for Research Assistance of the Minas Gerais State, (FAPEMIG-Brazil) [Grants CRA - RED-00060-23, and APQ-01942-22]. We also acknowledge research fellowships granted by CNPq to A.N-N. and W.L.A.

## Author Contributions

R.K.G.S., J.A.S., and W.L.A. designed the research; R.K.G.S., and J.A.S. performed most of the research with the support of W.B-S., M.F.S., T.W., J.R.V., G.V., R.R., and D.F.M.N.; A.A.A., C.R., A.R.F., and A.N-N. contributed new reagents/analytic tools; J.A.S., A.R.F., A.N-N., and W.L.A. analysed the data; and R.K.G.S., J.A.S., and W.L.A. wrote the article with input from all the others.

---

**Table S1.**
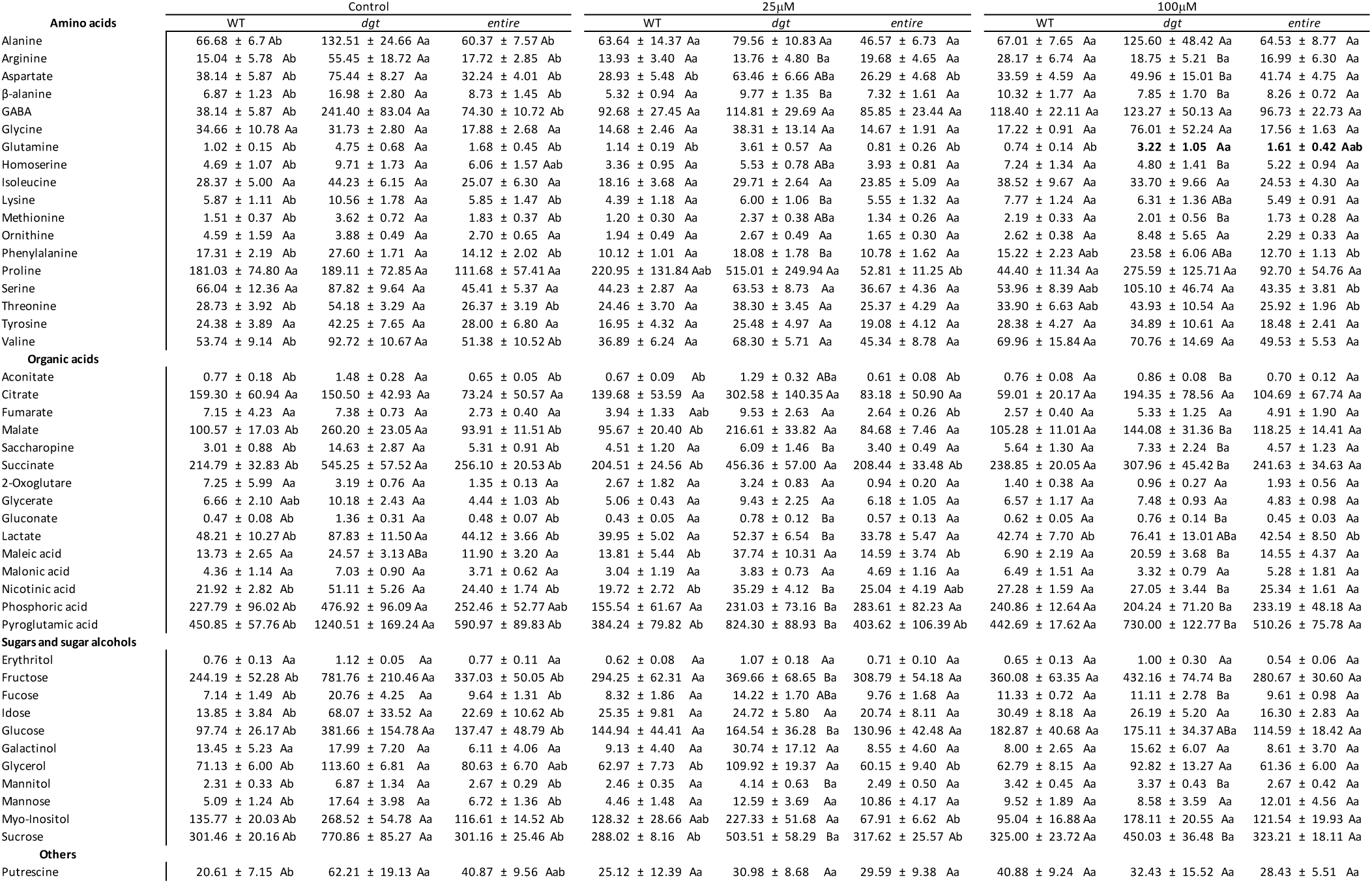
Metabolite profile of roots from mature tomato plants (35 DAG) with reduced auxin signalling (*diageotropica*, *dgt*) or enhanced auxin signalling (*entire*), compared to the wild-type (WT, *Solanum lycopersicum* cv. Micro-Tom) under Al³⁺ stress (25 μM and 100 μM). Values are expressed as mean ± SE (n = 6). Lowercase letters compare genotypes within the same treatment, while uppercase letters compare the same genotype under different treatments. Means followed by the same letters do not differ significantly according to Tukey’s test (*p* < 0.05).

